# Omicron-specific mRNA vaccine elicits potent immune responses in mice, hamsters, and nonhuman primates

**DOI:** 10.1101/2022.03.01.481391

**Authors:** Yi Wu, Yanqiong Shen, Namei Wu, Xinghai Zhang, Shaohong Chen, Chang Yang, Junhui Zhou, Yan Wu, Da Chen, Li Wang, Yuye Wang, Jiejie Xu, Ke Liu, Chao Wang, Huajun Zhang, Ninuo Xia, Sandra Chiu, Yucai Wang

## Abstract

SARS-CoV-2 has infected more than 400 million people around the globe and caused millions of deaths. Since its identification in November 2021, Omicron, a highly transmissible variant, has become the dominant variant in most countries. Omicron’s highly mutated spike protein, the main target of vaccine development, significantly compromises the immune protection from current vaccination. We develop an mRNA vaccine (S_Omicron_-6P) based on an Omicron-specific sequence. In mice, S_Omicron_-6P shows superior neutralizing antibodies inducing abilities to a clinically approved inactivated virus vaccine, a clinically approved protein subunit vaccine, and an mRNA vaccine (S_WT_-2P) with the same sequence of BNT162b2 RNA. Significantly, S_Omicron_-6P induces a 14.4∼27.7-fold and a 28.3∼50.3-fold increase of neutralizing activity against the pseudovirus of Omicron and authentic Omicron compared to S_WT_-2P, respectively. In addition, two doses S_Omicron_-6P significantly protects Syrian hamsters against challenge with SARS-CoV-2 Omicron variant and elicits high titers of nAbs in a dose-dependent manner in macaques. Our results suggest that S_Omicron_-6P offers advantages over current vaccines, and it will be helpful for those with weak immunity.

## INTRODUCTION

Severe acute respiratory syndrome coronavirus 2 (SARS-CoV-2) has infected more than 400 million people around the globe and caused several million deaths (Koh et al., 2021; Vogel et al., 2021). Since its discovery in November 2021, SARS-CoV-2 variant B.1.1.529, the World Health Organization (WHO) designation “Omicron”, has quickly spread and become dominant (Karim and Karim, 2021). The Omicron variant is highly transmissible and can infect human more quickly than other variants (Suzuki et al., 2022). Omicron currently represents ∼99% of the new infections in the US, Europe, and other major countries (www.gisaid.org/hcov19-variants/).

The Omicron variant carries approximately 30 mutations, some of which help it to escape the majority of existing SARS-CoV-2 neutralizing antibodies (nAbs) (Cao et al., 2021; Dejnirattisai et al., 2022a; Flemming, 2022; Planas et al., 2021). Most spike (S) protein monoclonal antibodies could no longer neutralize the Omicron variant. Convalescent individuals previously infected with other variants have little nAbs against Omicron and can be re-infected (Cele et al., 2021; Sun et al., 2022). Several studies show that the Omicron variant significantly weakened or knocked out the protection conferred by two vaccine doses. After a vaccine booster shot, vaccinees’ sera (post-vaccination sera) show enhanced nAb titers but are still around 20-fold less potent in neutralizing the Omicron variant than other variants (Hu et al., 2022; Liu et al., 2022; Planas et al., 2021; VanBlargan et al., 2022).

On the bright side, the third dose of current major vaccines significantly reduced the risk of hospitalization, severe illness, and death caused by Omicron. The Centers for Disease Control and Prevention (CDC) of America reported that a third vaccination prevented Omicron infected people from emergency room visits or urgent care with 82% and 90% effectiveness, respectively (Pia and Rowland-Jones, 2022; Thompson et al., 2021). However, for those with disadvantages, such as old age, pre-existing conditions, or being vaccinated with less potent vaccines, Omicron still poses a considerable threat. A recent phase 4 clinical trial in Brazil indicates a significant fraction of people who received three doses still have Omicron neutralization titers lower or barely above the limit to be considered seropositive (Malik et al., 2022; Mistry et al., 2021). Thus, an Omicron effective vaccine is urgently needed. Here we develop an Omicron variant sequence-based mRNA vaccine which is much more potent in inducing nAbs in multiple animal models against Omicron challenge than the original wild-type mRNA vaccine, inactivated virus vaccine, and protein subunit vaccine, and importantly, provides complete protection in hamster model at the dose as low as 1 μg.

## RESULTS

For full-length Omicron-specific mRNA vaccine design (named S_Omicron_-6P), we adopted the “hexapro” spike protein sequence as the backbone for its enhanced stability of prefusion conformation and substituted the respective sequences with the Omicron mutations (Table S1) (Hsieh et al., 2020). Modified Omicron mRNA was synthesized with high purity through in vitro transcription (Figure S1A). Robust expression of Omicron spike protein on HEK293T cell surface was detected after transfection with immunofluorescence (Figure S1B). The mRNA was then encapsulated into even-sized lipid nanoparticles (LNP) to generate the final vaccine product, S_Omicron_-6P, whose size is 110 nm on average (Figure S1C). We adopted the BNT162b2 RNA sequence with the two proline mutations as the control mRNA vaccine (S_WT_-2P) (Vogel et al., 2021).

We first tested the humoral responses to the immunogenicity of the vaccine. Mice were vaccinated twice and were sacrificed after two doses of S_Omicron_-6P (Figure 1A). We observed a significant increase in both total B cells (CD19^+^) and plasma B cells (CD138^+^B220^-^) in the spleens of S_Omicron_-6P immunized mice (Figure 1B and 1C), indicating S_Omicron_-6P can induce B cell responses. Then we performed a head-to-head comparison of S_Omicron_-6P *versus* S_WT_-2P, along with two clinically approved vaccines, one inactivated virus vaccine, and one protein subunit vaccine, on immunogenicity in BALB/c mice. Mice were vaccinated twice at various doses of each vaccine, and antibodies in the sera were measured one week after the second vaccination (Figure 1A). The antigen-specific IgG geometric mean titers (GMTs) were measured against Omicron Spike trimer protein with ELISA (Figure 1D and S2). Both S_Omicron_-6P and S_WT_-2P elicited IgG antibodies in a dose-dependent manner. At 5 and 10 µg dose levels, S_Omicron_-6P induced significantly higher IgG than S_WT_-2P, by 1.8-and 2.3-fold, respectively. The entry inhibition by serum of immunized mice was measured in a neutralization assay using vesicular stomatitis virus (VSV)-based Omicron pseudovirus. Dramatically but not surprisingly, the S_Omicron_-6P vaccinated mice elicited 14.4∼27.8-fold higher serum neutralizing activity than those by S_WT_-2P at all three dose groups (Figure 1E and S3-S4). Next, 50% virus-neutralization GMTs were measured by an Omicron-neutralization assay. As expected, 28.3∼50.3-fold higher neutralizing titers were observed from mice immunized with S_Omicron_-6P than those immunized with S_WT_-2P, using a plaque reduction neutralization test with authentic Omicron (Figure 1F). By contrast, two doses of immunization using inactivated virus vaccine or protein subunit vaccine hardly induced any Omicron nAbs in mice (Figure 1E-1F and S5-S6). These results suggest that S_Omicron_-6P is potent in inducing Omicron-specific antibodies.

**Figure 1.**
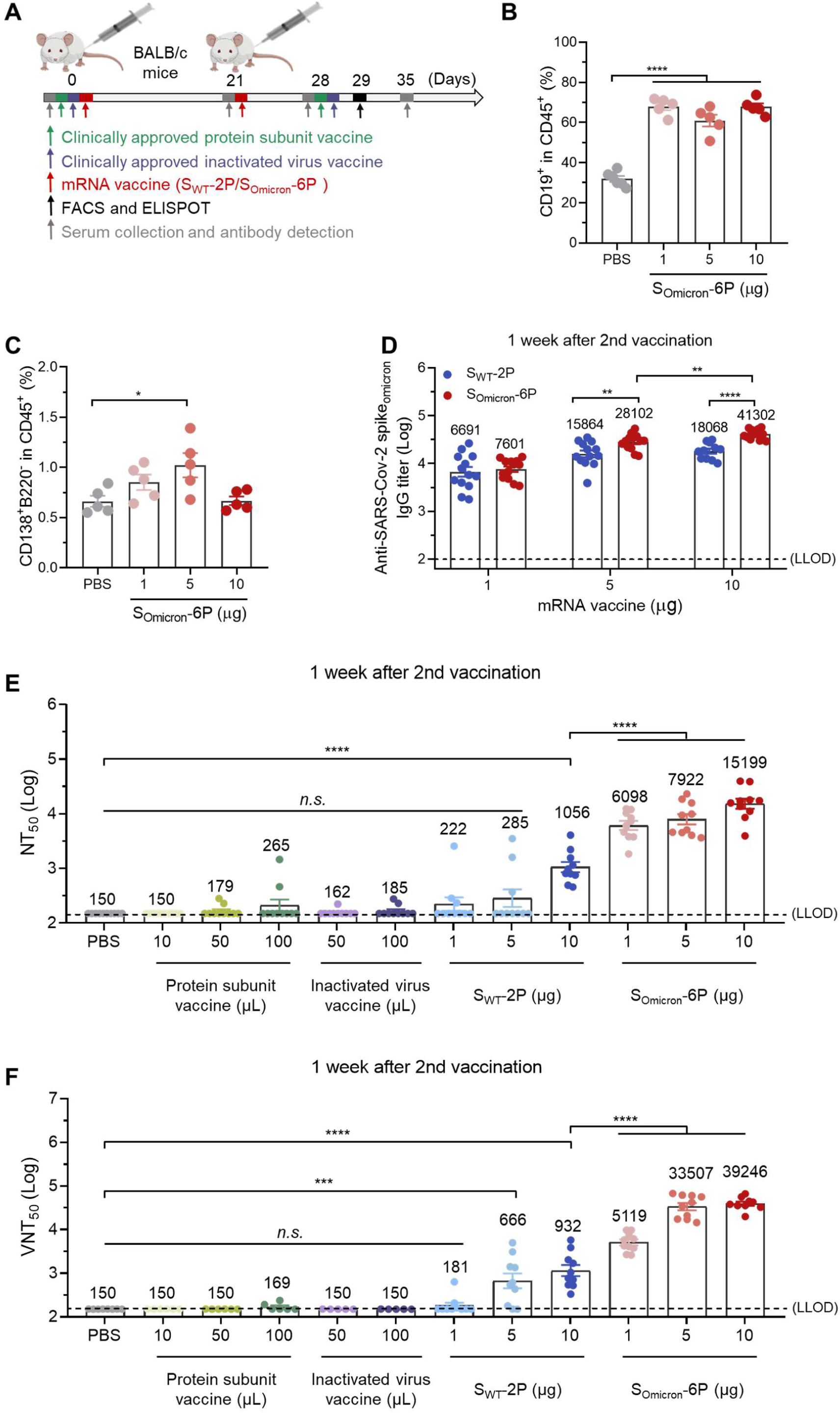
**S_Omicron_-6P Induces Antigen-Specific Humoral Immune Responses in Mice.** (A) Schematic diagram of immunization and sample collection schedule in mice. Female BALB/c mice were immunized on a two-dose schedule with S_WT_-2P, S_Omicron_-6P, protein subunit vaccine using a dimeric form of the receptor-binding domain of wild-type SARS-CoV-2, or inactivated vaccine of wild-type SARS-CoV-2. (B-C) Percentages of (B) B cells and (C) plasma cells in spleen after immunized with different doses of S_Omicron_-6P. (D) The Omicron SARS-CoV-2 variant specific IgG antibody titers were determined by ELISA (lower limit of detection (LLOD) = 100). (E) Neutralization titers (NT_50_) were determined by recombinant vesicular stomatitis virus (VSV)-based pseudovirus (Omicron variant) neutralization assay (LLOD = 150). (F) SARS-CoV-2 Omicron 50% virus-neutralization titers (VNT_50_) were determined by a plaque reduction neutralization test (LLOD = 150). Data are shown as mean ± SEM. Significance was calculated using one-way ANOVA with multiple comparisons tests (*n.s*., not significant, *p < 0.05, **p < 0.01, ***p < 0.001, ****p < 0.0001)

**Figure 2.**
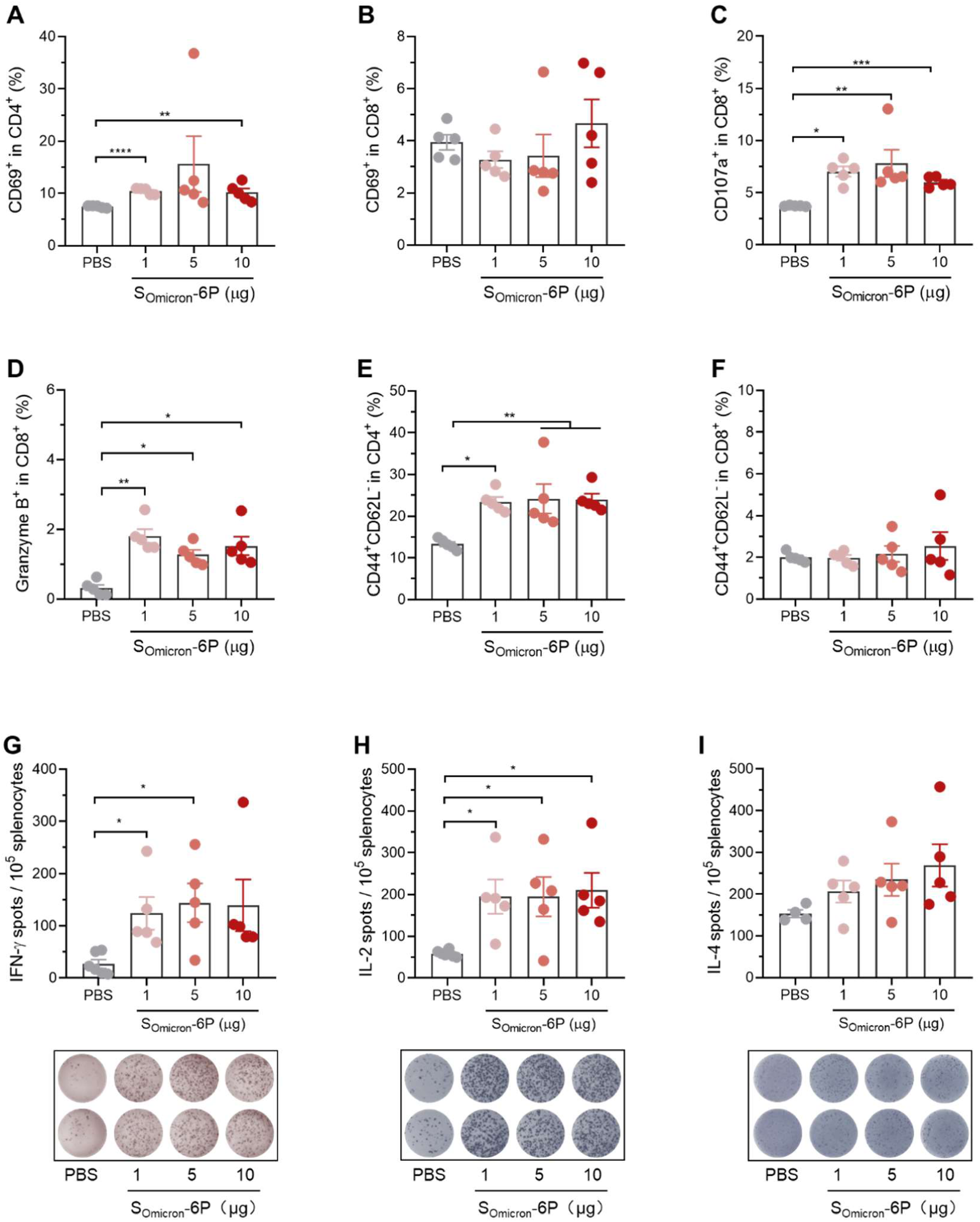
**S_Omicron_-6P Induces Antigen-Specific Celluar Immune Responses in Mice.** Female BALB/c mice were immunized with 0, 1, 5 or 10 µg S_Omicron_-6P. Twenty-nine days after the first immunization, mice were euthanized and their spleens were collected for T cell response and phenotyping analysis. (A-B) The percentages of activated (CD69^+^) (A) CD4^+^ and (B) CD8^+^ among CD4^+^ and CD8^+^ T cells. (C-D) The percentages of cytotoxic (CD107a^+^ and Granzyme B^+^) T cells among CD8^+^ T cells. (E-F) The percentages of effector memory (CD44^+^CD62L^-^) cells among (E) CD4^+^ and (F) CD8^+^ T cells. (G-I) ELISPOT assay for (G) IFN-γ, (H) IL-2, and (I) IL-4 in splenocytes. Splenocytes were harvested and re-stimulated with SARS-CoV-2 S protein peptide mix for 24 h on day 29 after first immunization. Data are shown as mean ± SEM. Significance was calculated using one-way ANOVA with multiple comparisons tests (*p < 0.05, **p < 0.01, ***p < 0.001, ****p < 0.0001)

**Figure 3.**
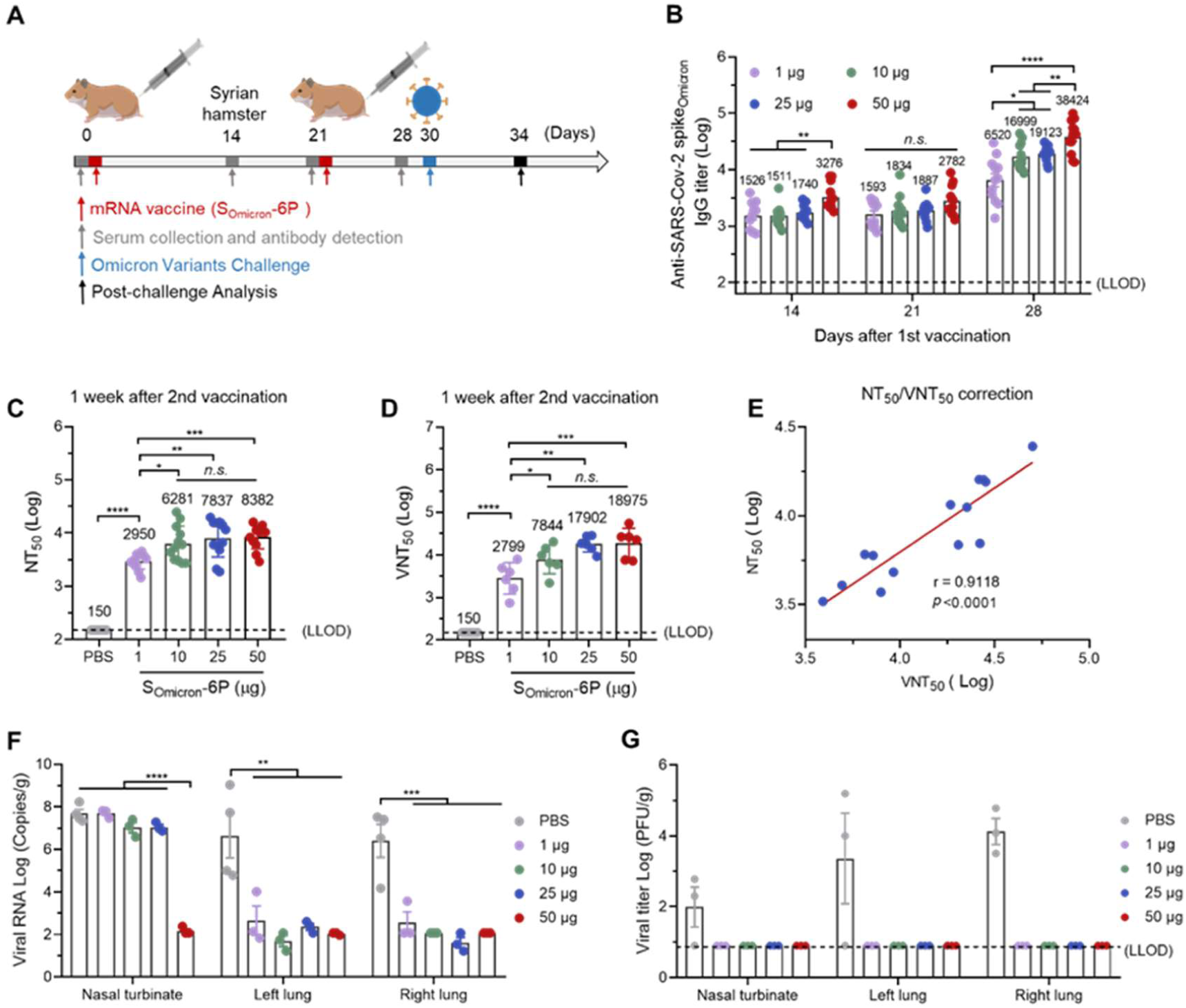
**S_Omicron_-6P Provides Robust Protection against Omicron in Syrian Hamsters.** (A) Schematic diagram of immunization and sample collection schedule in Syrian hamsters. Female hamsters were prime-vaccinated via the *i.m*. route on day 0 and boosted on day 21, with 0, 1, 10, 25, or 50 μg of S_Omicron_-6P. On day 30 after the initial immunization, hamsters were intranasally (*i.n.*) challenged with 1 × 10^4^ PFU of SARS-CoV-2 Omicron. On day 4 after infection, hamsters were euthanized for tissue collection. (B) The Omicron SARS-CoV-2 variant specific IgG antibody titers were determined by ELISA (lower limit of detection (LLOD) = 100). (C) NT_50_ values were determined by VSV-based pseudovirus (Omicron variant) neutralization assay (LLOD = 150). (D) VNT_50_ values were determined by a plaque reduction neutralization test (LLOD = 150). (E) Pearson correlation of VSV-SARS-CoV-2 (Omicron variant) VNT_50_ with live SARS-CoV-2 (Omicron variant) VNT_50_ for n = 14 random selected serum samples from mice immunized with S_Omicron_-6P. (F) Viral RNA load in the both lungs and nasal turbinates were determined by qRT-PCR. (G) Viral load expressed in PFU per gram of tissue in the both lungs and nasal turbinates at 4 dpi. Data are shown as mean ± SEM. Significance was calculated using one-way ANOVA with multiple comparisons tests (*n.s*., not significant, *p < 0.05, **p < 0.01, ***p < 0.001, ****p < 0.0001).

**Figure 4.**
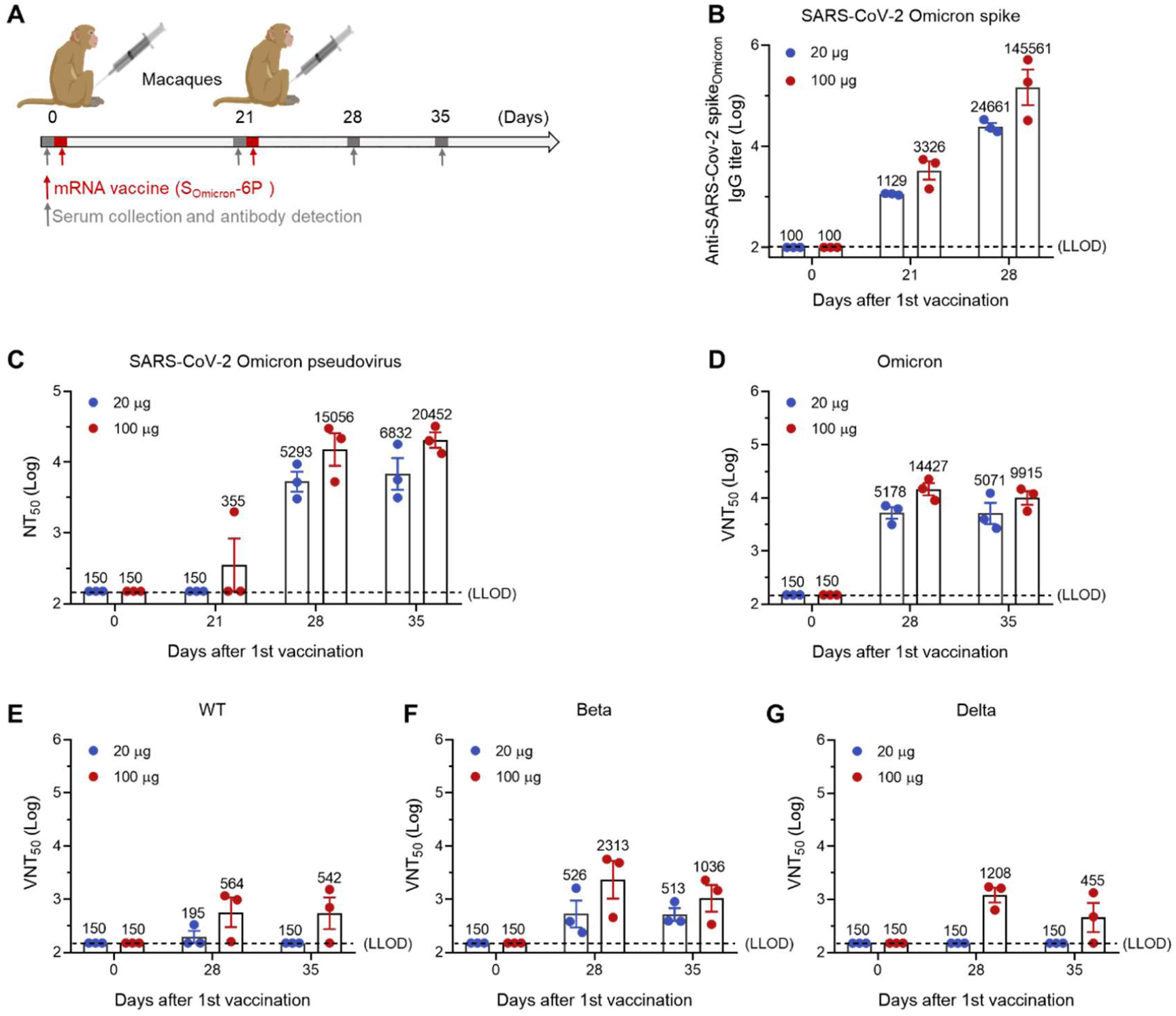
**S_Omicron_-6P Induces Omicron-Specific Immunity in Macaques.** (A) Study design. Male macaques were *i.m.* immunized with 20 or 100 μg S_Omicron_-6P and boosted with the same dose at 21-day interval. (B) The Omicron SARS-CoV-2 variant specific IgG antibody titers were determined by ELISA (lower limit of detection (LLOD) = 100). (C) NT_50_ values were determined by VSV-based pseudovirus (Omicron variant) neutralization assay (LLOD = 150). (D-G) VNT_50_ against (D) SARS-CoV-2 Omicron, (E) WT, (F) Beta, and (G) Delta that were determined by a plaque reduction neutralization test (LLOD = 150). Data are shown as mean ± SEM. Significance was calculated using one-way ANOVA with multiple comparisons tests.

We next investigated T cell responses in mice received two doses of S_Omicron_-6P. For T cell analysis in spleens, although the ratios of CD4^+^ and CD8^+^ T cells within CD45^+^ leukocytes remain unchanged, S_Omicron_-6P elicited significant increases in activated CD4^+^ (CD69^+^CD4^+^) and CD8^+^ (CD69^+^CD8^+^) T cells (Figure 2A, 2B, and Figure S7). We also noted that cytotoxic CD8^+^ T cells (CD107a^+^ and Granzyme B^+^), which play a crucial role in eliminating infected cells, have increased significantly after vaccination (Figure 2C and 2D). Furthermore, in the spleen of S_Omicron_-6P immunized mice, we observed extensive expansion of the effector memory CD4^+^ and CD8^+^ T cells (CD44^+^CD62L^-^), which mediate protective memory (Figure 2E and 2F).

Previous data indicate that mRNA vaccines induce T-helper-1 (Th1) -driven CD4^+^ T-cell responses (Laczko et al., 2020; Vogel et al., 2021). To investigate whether our S_Omicron_-6P activates an immune response similarly, we collected splenocytes from the immunized mice and re-stimulated them with the full-length S peptide mix. Using an enzyme-linked immunosorbent spot (ELISPOT) assay, we detected high levels of IFN-γ and interleukin-2 (IL-2) secreting Th1 cells in S_Omicron_-6P immunized mice (Figure 2G and 2H). Nevertheless, no significant difference in T-helper-2 (Th2) cytokines interleukin-4 (IL-4) secretion was observed between the vaccinated and control mice (Figure 2I). Intracellular-cytokine-staining flow cytometry had consistent results (Figure S8). These data confirmed that S_Omicron_-6P induces a Th1-biased immune response.

The Syrian hamster has been demonstrated as a suitable animal model for SARS-CoV-2 infection (Munoz-Fontela et al., 2020). Five groups of hamsters were vaccinated on day 0 and day 21 with either 1, 10, 25, and 50 µg of S_Omicron_-6P or PBS. The hamster sera were collected and evaluated for vaccine immunogenicity on day 14, 21 and 28 (Figure 3A). A significant amount of IgG against S protein was detected on day 14 and 21 after the first immunization, but no apparent dose-dependency was observed. However, the second dose boosts S antibodies more than ten times one week later (on day 28) (Figure 3B and S9). The pseudovirus assay showed that high neutralizing antibody titers were elicited even by 1 µg dose of S_Omicron_-6P (Figure 3C and S10). In line with this, high levels of neutralizing activity against authentic Omicron in S_Omicron_-6P vaccinated animals (Figure 3D). Moreover, we observed a strong correlation of pseudovirus assay with the authentic Omicron neutralization assay, with a correlation co-efficiency of 0.91 (Figure 3E).

On day 30, some hamsters were challenged with 1 × 10^4^ plaque-forming units (PFU) of authentic Omicron virus via intranasal route and sacrificed 4 days later. As shown in Figure 3F, only a trace amount of viral RNA was detected in the lung tissue of vaccinated animals with a little more for the 1 μg group, which is a 4-5 magnitude reduction than the control group. Infectious virus in lung tissue was determined with plaque assay, resulting in no detectable virus in both lungs and nasal turbinates of all vaccinated animals, including the lowest dose group, but while markedly levels of virus in the PBS group (Figure 3G). These data demonstrate that S_Omicron_-6P provides robust protection against the infection of Omicron.

The immunogenicity of S_Omicron_-6P was also evaluated in non-human primates. Macaques were immunized with either 20 or 100 µg of S_Omicron_-6P twice at a 21-day interval (Figure 4A). Similar to the results in mice and hamsters, the first immunization generates some level of IgG and almost no nAbs, with only one macaque producing little nAbs (Figure 4B-4C and S11-12). However, one week after the second immunization, all the macaques responded vigorously to the second immunization in one week and kept producing higher nAbs on day 35. Consistent with this, sera of the macaques showed vigorous neutralization activity against authentic Omicron on both day 28 and 35 (Figure 4D). Finally, we tested whether the nAbs elicited by S_Omicron_-6P could provide cross-protection against other SARS-CoV-2 variants. Although S_Omicron_-6P vaccinated macaques produced high nAbs against the Omicron variants, we detected relatively lower nAbs production against the wild-type, Beta, or Delta variants (Figure 4E-4G).

## DISCUSSION

Although the fast-spreading Omicron variant seems to cause less severe symptoms, the death toll keeps rising due to Omicron’s high transmissible ability (Garcia-Beltran et al., 2022). The original forms of mRNA vaccines, which have achieved remarkable clinical efficacy in protecting against prior variants of SARS-CoV-2, fail to provide as strong protection against Omicron as before. For example, the mRNA vaccine, BNT162b2, has over 90% efficacy against the WA1 strain; however, the efficacy dropped to around 30-50% for Omicron (Dejnirattisai et al., 2022b). The neutralizing antibody titers evoked by BNT162b2 dropped by 15-20 folds. Therefore, the fast evolution of the virus compelled us to develop Omicron-specific mRNA vaccines.

To generate an Omicron-specific mRNA vaccine, we introduced all the mutations within the S protein in the mRNA sequence and designed the sequence to express prefusion S protein *via* six consecutive proline substitutions (S_Omicron_-6P). S_Omicron_-6P induced a 14.4∼27.7-fold and a 28.3∼50.3-fold increase of neutralizing activity against the pseudovirus of Omicron and authentic Omicron compared to S_WT_-2P, respectively. Assuming that neutralizing antibody titers positively correlate to the protection efficacy, we anticipate that our Omicron-specific mRNA vaccine would restore its clinical effectiveness to at least 90%. Both the 6P design and the inclusion of all Omicron-specific mutations may contribute to the strong immunity evoked by the Omicron-specific mRNA vaccine.

Further immunology analysis indicates that the Omicron-specific mRNA vaccine induces multicomponent immune responses, including memory B and T cell responses, a Th1-biased T cell immunity, and a cytotoxic T cell response. The induction of immune response is similar to the prior version of mRNA vaccines, suggesting that changes in mRNA sequence do not alter the generic mechanisms of how mRNA-LNP-based vaccines activate cellular immunity.

In this study, we have conducted a comprehensive analysis of the effects of the Omicron-specific mRNA vaccine using several animal models, including mice, Syrian hamsters, and macaques. All the tested animals developed strong immune responses and were well protected from developing disease against Omicron virus challenge in hamsters. We noticed that in Syrian hamster model, only the highest dose of vaccines shows effective reduction of RNA copies in the nasal turbinate where SARS-COV2 replicates the most on day 4. In addition, no live Omicron virus was detected in nasal turbinates of all the S_Omicron_-6P-vaccinated hamsters. Note that we infected the animals by directly adding the Omicron viruses into the nasal cavity, and it may take a more extended period for the viral RNAs to degrade.

Most importantly, our data strongly suggest that two doses of Omicron-specific mRNA vaccine provide enhanced protection against Omicron compared with two doses of the prior version of mRNA vaccines with a booster. We note that several preprint studies published on bioRxiv indicate that the Omicron-specific booster offered no better protection against the Omicron variant than the S_WT_-2P booster (Ying et al., 2022). Note that the experimental settings of our study are different from these studies. We compare the immune protection effect against Omicron between Omicron-specific mRNA vaccines and wild type mRNA vaccines on naïve animals that have not been exposed to any of the prior vaccines. Our data provide strong evidence to show that Omicron-specific mRNA vaccines elicit enhanced protection for naïve animals. We think that highly the mutated S protein of Omicron has significantly compromised the immune memories induced by the prior version of vaccines so that a single dose of a booster, no matter Omicron-specific or not, is not potent enough to elicit immune protection effectively. Based on the results from our study, we urge that those with weaker immune systems should get at least two doses of Omicron-specific mRNA vaccines instead of getting a booster in addition to two doses of vaccines against prior SARS-CoV-2 variants. Furthermore, our data also suggest that Omicron-specific vaccines show considerable cross-protection against Beta variants, but lower protection against wild type and Delta variant. This also urges the necessity for development of multi-valent vaccine to fight against the evolution of SARS-CoV-2.

## ACKNOWLEDGMENTS

This work was supported by the National Key R&D Program of China (2020YFA0710700), the National Natural Science Foundation of China (52025036, 51961145109), the Fundamental Research Fund for the Central Universities (WK9100000014, WK2480000006), and the project of collaborative innovation for colleges of Anhui province (NO. GXXT-2021-070). This work was partially carried out at the USTC Center for Micro and Nanoscale Research and Fabrication. We thank Jia Wu, Jun Liu and Hao Tang from Wuhan Institute of Virology for managing of BSL-3 facility, where all the authentic SARS-CoV-2 experiments were conducted. We also thank National Virus Resource Center for providing the Omicron variant (CCPM-B-V-049-2112-18). We thank Weiheng Chen from the Animal Facility of USTC, where the mice, hamsters, and macaques were vaccinated. We thank the Joint Laboratory of Innovation in Life Sciences University of Science and Technology of China (USTC) and Changchun Zhuoyi Biological Co. Ltd.

## AUTHOR CONTRIBUTIONS

C. W., N.-N.X., Y.-C.W., and S.C. supervised the project. Yi W., Y.-Q.S., N.-M.W., Y.- C.W., and S.C. conceived the experiments. Yi W., Y.-Q.S., N.-M.W., X.-H.Z., S.-H.C., C.Y., H.-J.Z., Yan W., D.C., L.W., Y.-Y.W., J.-J.X., K.L. conducted the experiments and analyzed the data. Yi W., Y.-Q.S., N.-M.W., N.-N.X., and Y.-C.W. wrote and revised the manuscript. C. W., H.-J.Z., and S.C. revised the manuscript. Y.-Y.W., J.-J.X., and K.L. are employees of Hefei RNAlfa Biotech. All authors read and approved the manuscript.

## DECLARATION OF INTERESTS

N.-N.X., Y.-C.W. are co-inventors on pending patent applications related to the Omicron mRNA vaccine. The other authors declare no known competing financial interests or personal relationships that could have appeared to influence the work reported in this paper.

## KEY RESOURCES TABLE

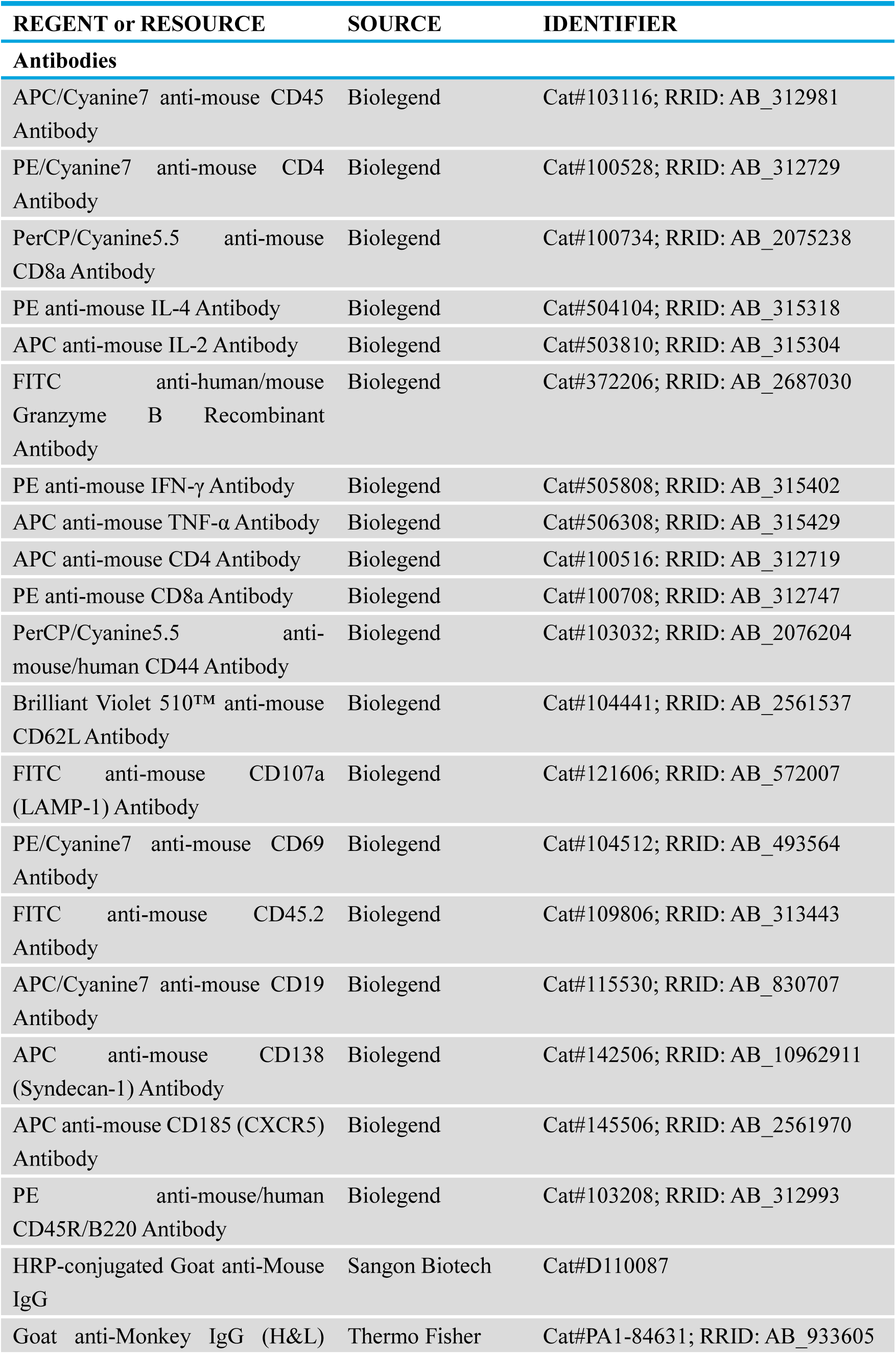
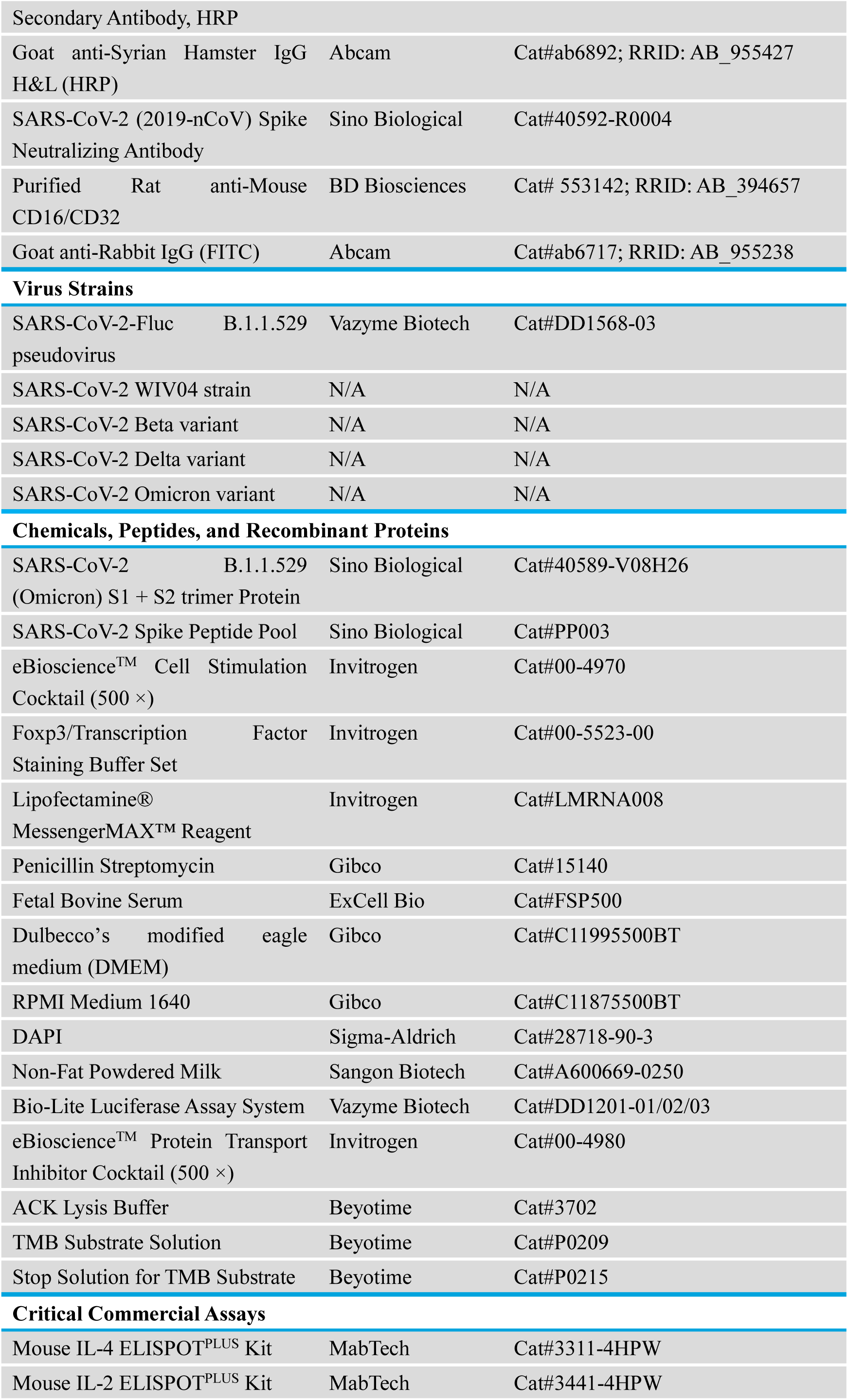
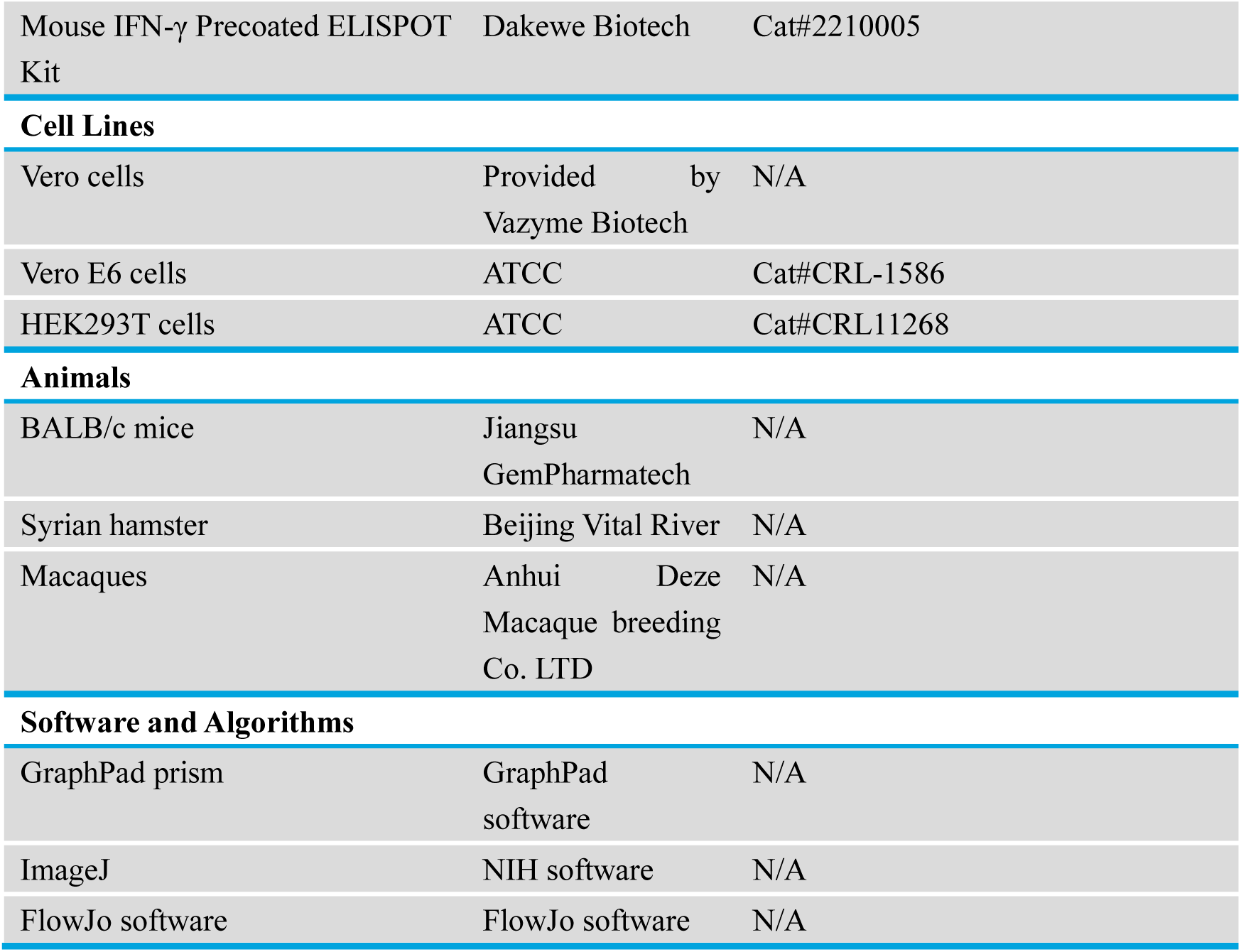

## EXPERIMENTAL MODEL AND SUBJECT DETAILS

### Ethics statement

All mouse and macaque studies were conducted under protocols approved by the Institutional Animal Care and Use Committee of the University of Science and Technology of China. All procedures performed on Syrian hamster were in accordance with regulations and established guidelines, and were reviewed and approved by the Animal Ethics Committee of the Wuhan Institute of Biological Products (WIBP) (WIBP-AII382020001). The animals received care in compliance with the guidelines outlined in the Guide for the Care and Use of Laboratory Animals.

### Cells and viruses

HEK293T cells, Vero cells, and Vero E6 cells were cultured in Dulbecco’s modified Eagle’s medium (DMEM, Gibco) supplemented with 10% fetal bovine serum (FBS, ExCell Bio) and 1% penicillin-streptomycin (Gibco) at 37°C under a 5% CO_2_ atmosphere. The SARS-CoV-2 WIV04 strain was initially isolated from a COVID-19 patient in 2019 (GISAID, accession no. EPI_ISL_402124); Beta variant (NPRC2.062100001) was kindly provided by Chinese Center for Disease Control and Prevention, and Delta variant (B.1.617.2; GWHBEBW01000000) by Prof. Hongping Wei; Omicron variant was isolated from a throat swab of a patient from Hong Kong by the Institute of Laboratory Animal Sciences, Chinese Academy of Medical Sciences (CCPM-B-V-049-2112-18). All processes in this study involving authentic SARS-CoV-2 were performed in a BSL-3 facility.

## METHOD DETAILS

### mRNA design and synthesis

Spike (S) protein encoded by S_WT_-2P vaccine was designed from original ancestral SARS-CoV-2 WA1 (GenBank MN908947.3), S_Omicron_-6P was based on a background of S sequences from SARS-CoV-2 variant Omicron (B.1.1.529) (GISAID: GR/484A). The template for the S_WT_-2P mRNA is a DNA fragment encoding SARS-CoV-2 S with K986P and V987P substitutions. The template for the S_Omicron_-6P mRNA is a DNA fragment encoding Omicron variant S with F817P, A892P, A899P, A942P, K986P, and V987P substitutions. Both mRNAs were synthesized in vitro using an optimized T7 RNA polymerase-mediated transcription reaction with complete replacement of uridine by N1-methyl-pseudouridine. The reaction included a DNA template containing the open reading frame flanked by 5’ untranslated region (UTR) and 3’ UTR sequences and was terminated by an encoded poly A tail.

The mRNA was purified by oligo-dT affinity purification, buffer exchanged by tangential flow filtration into sodium acetate, and sterile filtered. RNA integrity was assessed by microfluidic capillary electrophoresis (Fragment Analyzer systems 5200, Agilent), and the concentration, pH, residual DNA, proteins, and dsRNA impurities of the solution were determined. The mRNA 5’ capping efficiency and 3’-polyadenosine (poly A) tail of mRNAs was studied using liquid chromatography coupled to mass spectrometry (LC-MS).

### mRNA vaccine production

mRNAs were encapsulated in LNPs using a modified procedure of a method previously as previously described (Maier et al., 2013) wherein an ethanolic lipid mixture of ionizable cationic lipid, phosphatidylcholine, cholesterol, and polyethylene glycol-lipid was rapidly mixed with an aqueous solution containing mRNA. The drug product underwent analytical characterization, which included the determination of particle size and polydispersity, encapsulation, pH, endotoxin, and bioburden, and the material was deemed acceptable for in vivo study.

### mRNA transfection

HEK293T were seeded in 24-well plates at 1.5 × 10^4^ cells/well. After 12 h, the cells were transfected with S_Omicron_-6P mRNA using Lipofectamine^®^ Messenger MAX™ Reagent (Invitrogen). And 6 h later, the medium was replaced with DMEM medium (Gibco).

### Vaccine antigen detection by immunofluorescence

Transfected HEK293T cells were fixed in 4% paraformaldehyde (PFA) and permeabilized in PBS/0.1% Triton X-100. Free binding sites were blocked with 1% BSA for 0.5 h at room temperature. Then cells were incubated with SARS-CoV-2 S neutralizing antibody (Sino Biological, 40592-R0004) that recognizes Omicron S protein. The cells were stained with an anti-rabbit fluorescent IgG secondary antibody, and nucleus DNA was stained with DAPI (Sigma-Aldrich). Images were acquired with a laser scanning confocal microscope (Nikon A1).

### Mouse immunizations

Female BALB/c mice (8–12 weeks old) were randomly allocated to groups. For S_WT_-2P and S_Omicron_-6P mRNA groups, the mice were immunized intramuscularly with 1, 5, and 10 μg of mRNA vaccine, respectively. For clinically approved inactivated vaccines, the mice were intramuscularly (*i.m.*) immunized with 50 µL and 100 µL of the vaccine (500 µL/vial for an adult), respectively. For clinically approved protein subunit vaccine, mice were *i.m.* immunized with 10, 50, and 100 µL of the vaccine (500 µL/vial for an adult), respectively. Mice were immunized with the same dose at 21-day intervals for mRNA vaccines or 28-day intervals for inactivated vaccine or protein subunit vaccine. Sera were collected on day 0, 21, 28, and 35 after the first immunization to detect SARS-CoV-2 Omicron variant-specific IgG and nAbs titers as described below. Spleens of mice receiving different vaccines were collected on day 29 post the first immunization to evaluate immune responses by ELISPOT and flow cytometry as described below.

### Hamster immunization and challenge experiments

Four groups of female Syrian hamsters (6–10 weeks old) were vaccinated with 1, 10, 25, 50 μg of S_Omicron_-6P for prime-boost vaccine regimens. PBS *i.m.* immunization served as control. Formulations were administered by intramuscular injection to each hind leg. On day 21, all groups received their second vaccine dose. On day 30, groups of 5 female Syrian hamsters were challenged *i.n.* with 1 × 10^4^ PFU of the Omicron variants per animal after anesthetization with isoflurane. The SARS-CoV-2 Omicron virus was isolated from a patient’s throat swab of from Hong Kong by the Institute of Laboratory Animal Sciences, Chinese Academy of Medical Sciences (CCPM-B-V-049-2112-18). All processes in this study involving authentic SARS-CoV-2 were performed in a BSL-3 facility. Throughout the study, hamsters were monitored daily for weight changes.

### Macaque immunizations

Male macaques (3–5 years old) were randomly assigned to receive S_Omicron_-6P (20 or 100 µg) on day 0 (the day for the first vaccination) and 21. The vaccine was administered *i.m.* injection in the quadriceps muscle. Blood was collected on day 0, 21, 28, and 35 after the first immunization to detect Omicron variant S protein-specific IgG and nAbs as described below.

### Enzyme linked immunosorbent assay (ELISA)

Nunc Maxisorp ELISA plates (ThermoFisher) were coated with 100 ng per well of SARS-CoV-2 B.1.1.529 (Omicron) S1 + S2 trimer protein (Sino Biological) in PBS overnight at 4℃. The coated plates were washed 4 times with PBS and blocked with 5% skim milk powder in PBST (0.1% Tween-20 in PBS) for 2 h. After blocks, plates were incubated with serial dilutions of heat-inactivated sera in blocking buffer for 1 h at room temperature, followed by 4 washes. HRP-conjugated secondary antibody was diluted 1:10,000 in blocking buffer and incubated for 1 hour, followed by 4 washes. TMB (Beyotime) substrate was added and reacted under dark for 8 minutes. The absorbance was measured at 450 nm using a SpectraMax iD5 microplate reader.

### Pseudovirus neutralization assays

A recombinant vesicular stomatitis virus (VSV)-based pseudovirus neutralization assay was used to measure neutralizing antibodies. The SARS-CoV-2-Fluc B.1.1.529 pseudovirus (Vazyme Biotech, DD1568-03) was used. In brief, pseudovirus carrying a luciferase reporter and encapsulated in Omicron variant S proteins were incubated with six 4-fold serial dilutions of the heat-inactivated serum samples by DMEM (Gibco) for 1 h at 37℃. The mixture was then added to the Vero cells culture (Vazyme Biotech) in 96-well plates with DMEM /10% FBS/1% penicillin-streptomycin and incubated in a humidified cell culture chamber at 37°C with 5% CO_2_ for 24 hours. The medium was removed at the end of incubation, and 100 μL one-step luciferase detection reagent (Vazyme Biotech, DD1201-03) was added to each well. Luminescence in relative light units (RLUs) was measured by a luminometer (SpectraMax iD5, Molecular Devices) after 3 minutes of incubation at room temperature. Serum samples may be diluted to meet the initial volume requirement. RLUs of sample wells were normalized with positive control wells, and NT_50_ was calculated as EC50 by a normalized four-parameter sigmoid curve fit with constrains of EC50 > 0 and hillslope > 0 in Prism 8.0 (GraphPad).

### Plaque reduction neutralization assay

The plaque reduction neutralization assay was carried out as described before (Wang et al., 2020). Sera were inactivated at 56°C for 30 min before use. The sera were diluted 150-fold first, and then 3-fold serial dilutions were prepared in the maintenance medium. The virus suspension (0.25 mL, 600 PFU/mL) was mixed with an equal volume of antiserum at desirable dilution and incubated for 1 h. The mixture was added to monolayer cells in 24-well plates and incubated for 1 h. After removing of the mixture, 2 mL of maintenance medium containing 0.9% of methylcellulose were added to each well. The plates were incubated in a 5% CO_2_-air incubator at 37°C for 3–4 days. The neutralizing titer was calculated as reciprocal of the highest sera dilution suppressing 50% of plaque forming. Plaque reduction nAb titer (VNT_50_, 95% CI, challenge viruses used: 30–300 PFU/well) was calculated as the “inhibitor vs normalized response (Variable slope)” model in the GraphPad Prism 8.0 software.

### Flow cytometry

Sample processing: spleens were collected in PBS and homogenized through a 70 µm cell strainer using the stern end of a syringe plunger. Splenocytes were incubated in ACK lysis buffer to remove red blood cells, then passed through a 40 µm strainer to obtain a single-cell suspension.

Cell activation analysis: after preparing spleen single-cell suspensions, cells were immediately analyzed for activation markers. Cells were blocked by Fc-receptor blockade with anti-CD16/CD32 (BD Biosciences), and then stained for 30 minutes at 4℃ with the following antibody panel each diluted in PBS: APC/Cyanine7 anti-mouse CD45 antibody (Biolegend) or FITC anti-mouse CD45.2 antibody (Biolegend), APC anti-mouse CD4 antibody (Biolegend), APC/Cyanine7 anti-mouse CD19 antibody (Biolegend), PE anti-mouse CD8a antibody (Biolegend) or PE anti-mouse/human CD45R/B220 antibody (Biolegend), PerCP/Cyanine5.5 anti-mouse/human CD44 antibody (Biolegend), APC anti-mouse CD138 (Syndecan-1) antibody (Biolegend), Brilliant Violet 510™ anti-mouse CD62L antibody (Biolegend), FITC anti-mouse CD107a (LAMP-1) antibody (Biolegend), APC/Cyanine7 anti-mouse CD19 antibody (Biolegend). Samples were analyzed on the CytoFLEX LX flow cytometer (Beckman Coulter).

Intracellular cytokine staining: to measure antigen-specific T cells, spleen cells were stimulated with the full-length S peptide mix (Sino Biological) and eBioscience^TM^ protein transport inhibitor cocktail (Invitrogen) at 37℃, 5% CO_2_. RPMI medium 1640 served as a negative control and the combination of eBioscience^TM^ cell stimulation cocktail (Invitrogen) and eBioscience^TM^ protein transport inhibitor cocktail (Invitrogen) served as a positive control. Then samples were blocked by anti-CD16/CD32 blockade as above, stained for 30 minutes at 4℃ with the following antibody: APC/Cyanine7 anti-mouse CD45 antibody (Biolegend), PE/Cyanine7 anti-mouse CD4 antibody (Biolegend), PerCP/Cyanine5.5 anti-mouse CD8a antibody (Biolegend). Cells were washed and fixed and permeabilized using Foxp3/Transcription factor staining buffer set (Invitrogen), and stained intracellularly for 30 minutes in PBS with antibodies including: PE anti-mouse IL-4 antibody (Biolegend) or PE anti-mouse IFN-γ antibody (Biolegend), APC anti-mouse IL-2 antibody (Biolegend) and FITC anti-human/mouse Granzyme B recombinant antibody (Biolegend). Samples were analyzed on the CytoFLEX LX flow cytometer (Beckman Coulter).

### ELISPOT

IL-4 and IL-2 ELISPOT assays were performed with mouse IL-4 ELISPOT^PLUS^ kits and mouse IL-2 ELISPOT^PLUS^ kits according to the manufacturer’s instructions (Mabtech). IFN-γ ELISPOT assays were performed with mouse IFN-γ Precoated ELISPOT kits according to the manufacturer’s instructions (Dakewe Biotech). Briefly, a total of 5 × 10^5^ splenocytes in a volume of 200 µL was stimulated with the full-length S peptide mix (Sino Biological) (0.1 μg/mL final concentration per peptide). After incubation at 37℃, 5% CO_2_ for 18 h, the plates were washed, and biotinylated anti-mouse IFN-γ, IL-2, or IL-4 antibody was added to each well, following incubation of detection second antibodies. The air-dried plates were read using the automated ELISPOT reader (Mabtech IRIS FluoroSpot/ELISpot reader) for calculating spot-forming cells.

### Analysis of viral load by RT-qPCR

Viral RNA in lung tissues from challenged hamsters was quantified by one-step real-time RT-PCR as described before (Feng et al., 2020). Briefly, viral RNA was purified using the QIAamp Viral RNA Mini Kit (Qiagen), and quantified with HiScript® II One Step qRT-PCR SYBR® Green Kit (Vazyme Biotech) with the primers ORF1ab-F (5’-CCCTGTGGGTTTTACACTTAA-3’) and ORF1ab-R (5’-ACGATTGTGCATCAGCTGA-3’). The amplification procedure was set up as: 50°C for 3 min, 95°C for 30 s followed by 40 cycles consisting of 95°C for 10 s, 60°C for 30 s.

### Analysis of viral load by plaque assay

Virus titer was determined with plaque assay as previously described with slight modification (Zhang et al., 2020). Briefly, virus samples were serially 10-fold diluted with DMEM with 2.5% FBS, and inoculated to Vero cells or Vero E6 seeded overnight at 1.5 × 10^5^ /well in 24-well plates; after incubated at 37°C for 1 h, the inoculate was replaced with DMEM containing 2.5% FBS and 0.9% carboxymethyl-cellulose. The plates were fixed with 8% paraformaldehyde and stained with 0.5% crystal violet 3 days later. Virus titer was calculated with the dilution gradient with 10∼100 plaques. Plaque assays were performed in a BSL3 facility with strict adherence to institutional regulations.

## QUANTIFICATION AND STATISTICAL ANALYSIS

All data were analyzed with GraphPad Prism 8.0 software. No statistical methods were used to predetermine sample size, unless indicated. Unless specified, data are presented as mean ± SEM in all experiments. Analysis of variance (ANOVA) or t-test was used to determine statistical significance among different groups (*p < 0.05; **p < 0.01; ***p < 0.001; ****p < 0.0001).

## SUPPLYMENTRY TABLES AND FIGURES

**Table S1.** Amino Acid **Sequence Alignment of the Full S Protein of S_Omicron_-6P.** MFVFLVLLPLVSSQCVNLTTRTQLPPAYTNSFTRGVYYPDKVFRSSVLHSTQDLFLPFFSNVTWFHVISGTNGTKRFDNPVLPFNDGVYFASIEKSNIIRGWIFGTTLDSKTQSLLIVNNATNVVIKVCEFQFCNDPFLDHKNNKSWMESEFRVYSSANNCTFEYVSQPFLMDLEGKQGNFKNLREFVFKNIDGYFKIYSKHTPIIVRDLPQGFSALEPLVDLPIGINITRFQTLLALHRSYLTPGDSSSGWTAGAAAYYVGYLQPRTFLLKYNENGTITDAVDCALDPLSETKCTLKSFTVEKGIYQTSNFRVQPTESIVRFPNITNLCPFDEVFNATRFASVYAWNRKRISNCVADYSVLYNLAPFFTFKCYGVSPTKLNDLCFTNVYADSFVIRGDEVRQIAPGQTGNIADYNYKLPDDFTGCVIAWNSNKLDSKVSGNYNYLYRLFRKSNLKPFERDISTEIYQAGNKPCNGVAGFNCYFPLRSYSFRPTYGVGHQPYRVVVLSFELLHAPATVCGPKKSTNLVKNKCVNFNFNGLKGTGVLTESNKKFLPFQQFGRDIADTTDAVRDPQTLEILDITPCSFGGVSVITPGTNTSNQVAVLYQGVNCTEVPVAIHADQLTPTWRVYSTGSNVFQTRAGCLIGAEYVNNSYECDIPIGAGICASYQTQTKSHRRARSVASQSIIAYTMSLGAENSVAYSNNSIAIPTNFTISVTTEILPVSMTKTSVDCTMYICGDSTECSNLLLQYGSFCTQLKRALTGIAVEQDKNTQEVFAQVKQIYKTPPIKYFGGFNFSQILPDPSKPSKRSPIEDLLFNKVTLADAGFIKQYGDCLGDIAARDLICAQKFKGLTVLPPLLTDEMIAQYTSALLAGTITSGWTFGAGPALQIPFPMQMAYRFNGIGVTQNVLYENQKLIANQFNSAIGKIQDSLSSTPSALGKLQDVVNHNAQALNTLVKQLSSKFGAISSVLNDIFSRLDPPEAEVQIDRLITGRLQSLQTYVTQQLIRAAEIRASANLAATKMSECVLGQSKRVDFCGKGYHLMSFPQSAPHGVVFLHVTYVPAQEKNFTTAPAICHDGKAHFPREGVFVSNGTHWFVTQRNFYEPQIITTDNTFVSGNCDVVIGIVNNTVYDPLQPELDSFKEELDKYFKNHTSPDVDLGDISGINASVVNIQKEIDRLNEVAKNLNESLIDLQELGKYEQYIKWPWYIWLGFIAGLIAIVMVTIMLCCMTSCCSCLKGCCSCGSCCKFDEDDSEPVLKGVKLHYT

**Figure S1.**
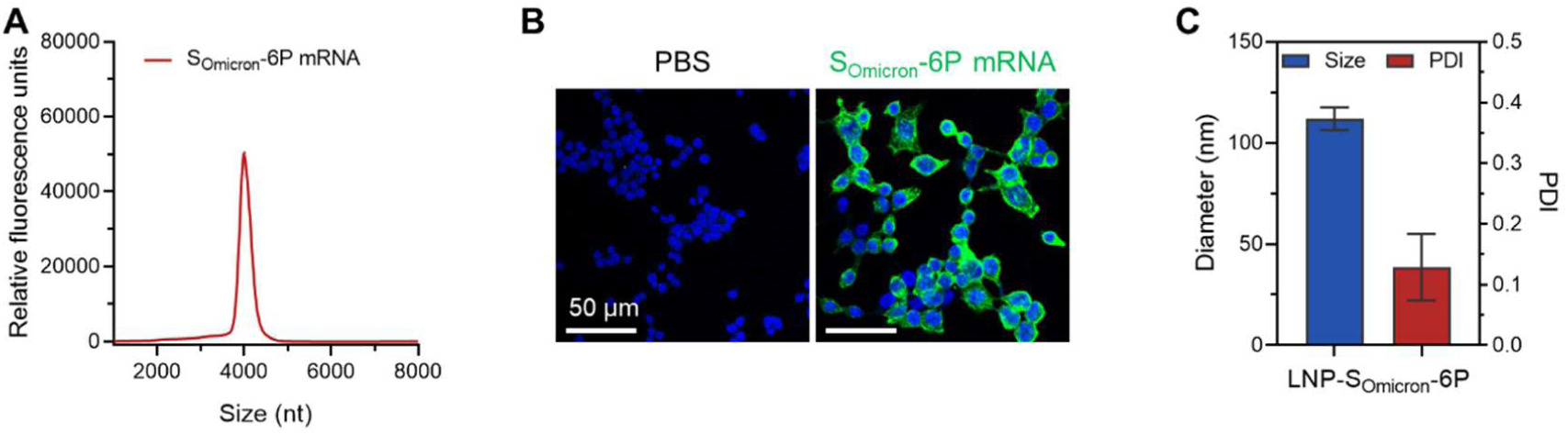
SARS-CoV-2 Omicron mRNA Vaccine Design and Characterization, Related to Figures 1-4. (A) Liquid capillary electropherograms of in vitro-transcribed S_Omicron_-6P mRNA. Peaks represent individual samples merged into one graph. (B) Immunofluorescence analysis of the expression of Omicron spike protein in HEK293T cells. (C) Sizes and polydispersity index (PDI) values of lipid nanoparticles (LNP) encapsulated with S_Omicron_-6P mRNA. Data are shown as mean ± SD.

**Figure S2.**
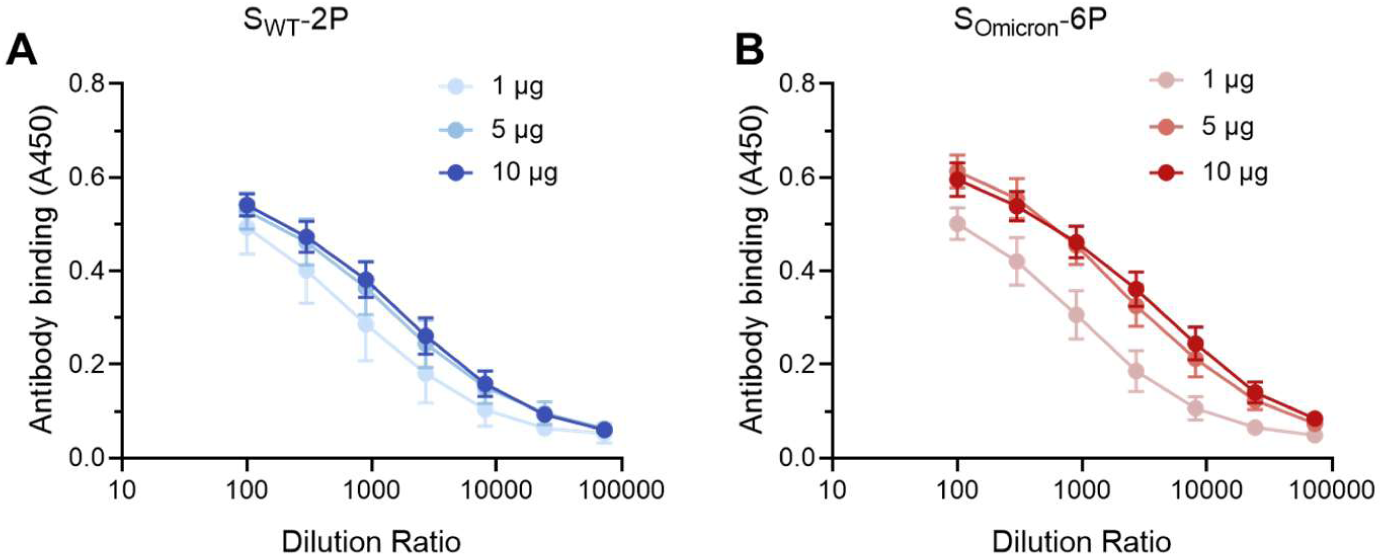
S_Omicron_-6P or S_WT_-2P Elicited Binding Antibodies in Mice, Related to Figure 1. (A-B) ELISA binding curves of (A) S_WT_-2P or (B) S_Omicron_-6P induced binding antibodies in mouse sera.

**Figure S3.**
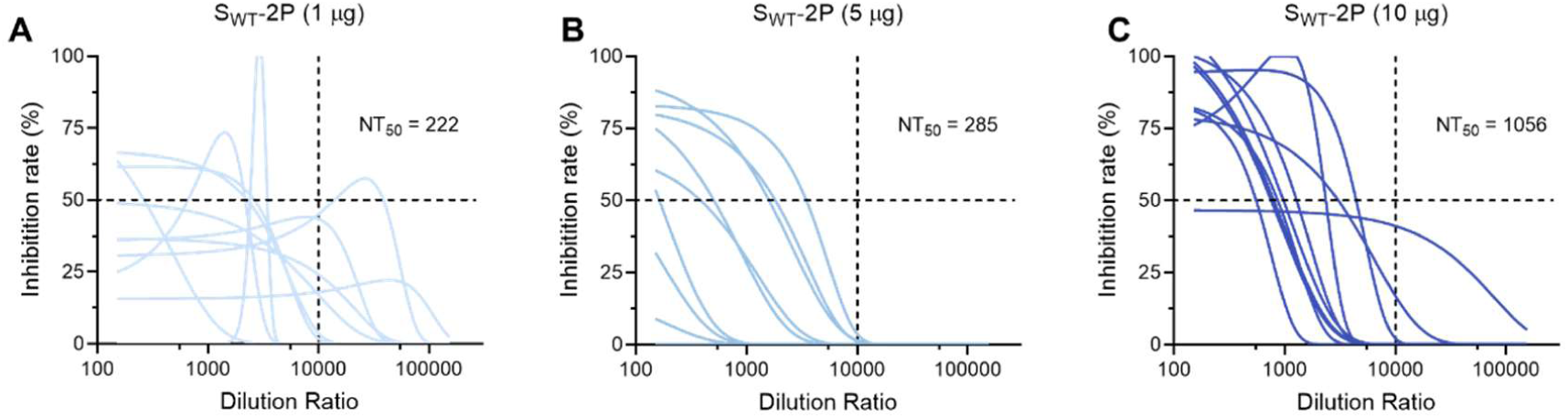
S_WT_-2P Induced Low Levels of nAbs Against SARS-CoV-2 Omicron Variant in Mice, Related to Figure 1. (A-C) Neutralization curves of (A) 1, (B) 5, and (C) 10 µg S_WT_-2P induced antibodies against pseudotyped and replication-deficient SARS-CoV-2 Omicron.

**Figure S4.**
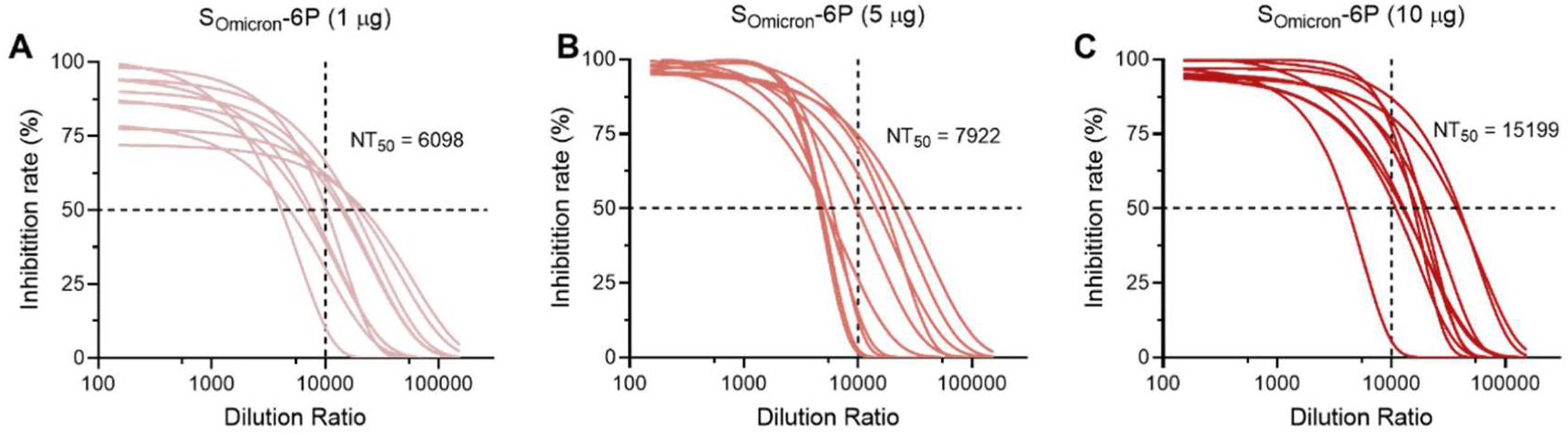
S_Omicron_-6P Induced High Levels of nAbs Against SARS-CoV-2 Omicron Variant in Mice, Related to Figure 1. (A-C) Neutralization curves of (A) 1, (B) 5, and (C) 10 μg S_Omicron_-6P induced antibodies against pseudotyped and replication-deficient SARS-CoV-2 Omicron.

**Figure S5.**
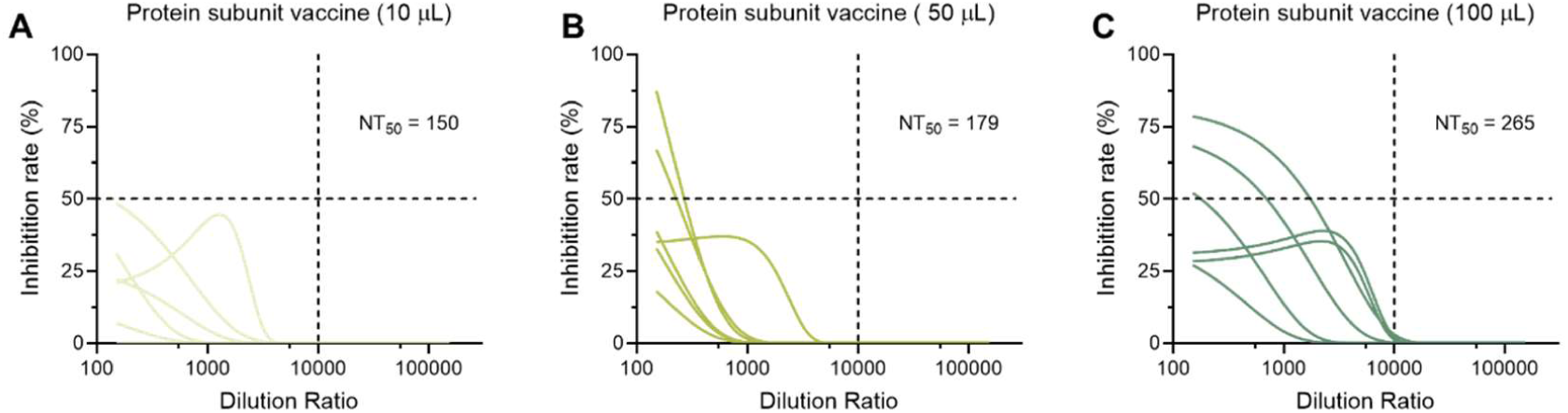
Clinically Approved Protein Subunit Vaccine Rarely Induced nAbs Against SARS-CoV-2 Omicron Variant in Mice, Related to Figure 1. (A-C) Neutralization curves of (A) 10, (B) 50, and (C) 100 μL protein subunit vaccine (500 µL/vial for an adult) induced antibodies against pseudotyped and replication-deficient SARS-CoV-2 Omicron.

**Figure S6.**
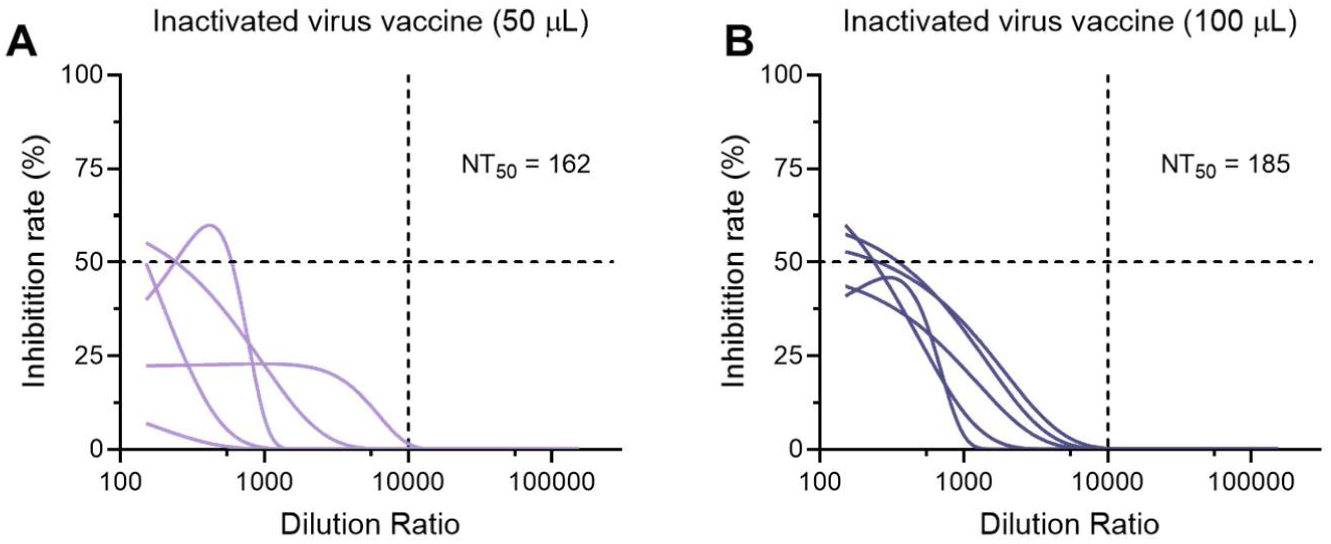
Clinically Approved Inactivated Virus Vaccine Rarely Induced nAbs Against SARS-CoV-2 Omicron Variant in Mice, Related to Figure 1. (A-B) Neutralization curves of (A) 50, and (B) 100 μL inactivated virus vaccine (500 µL/vial for an adult) induced antibodies against pseudotyped and replication-deficient SARS-CoV-2 Omicron.

**Figure S7.**
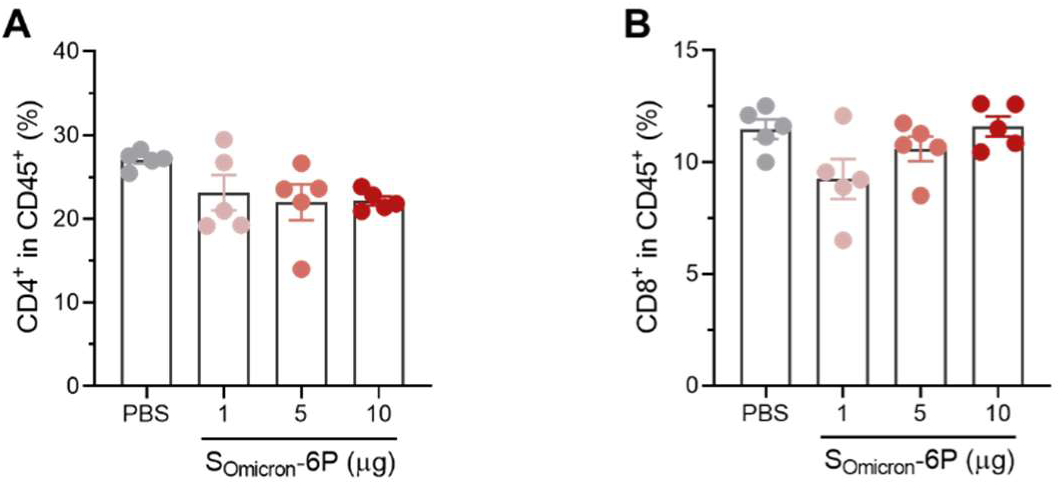
The Percentages of CD4+ and CD8+ T Cells Among Lymphocytes in Spleen, Related to Figure 2. Female BALB/c mice were immunized with 0, 1, 5 or 10 µg S_Omicron_-6P. Twenty-nine days after the first immunization, mice were euthanized and their spleens were collected for T cell response and phenotyping analysis. (A-B) The percentages of (A) CD4^+^ and (B) CD8^+^ T cells among lymphocytes in spleen.

**Figure S8.**
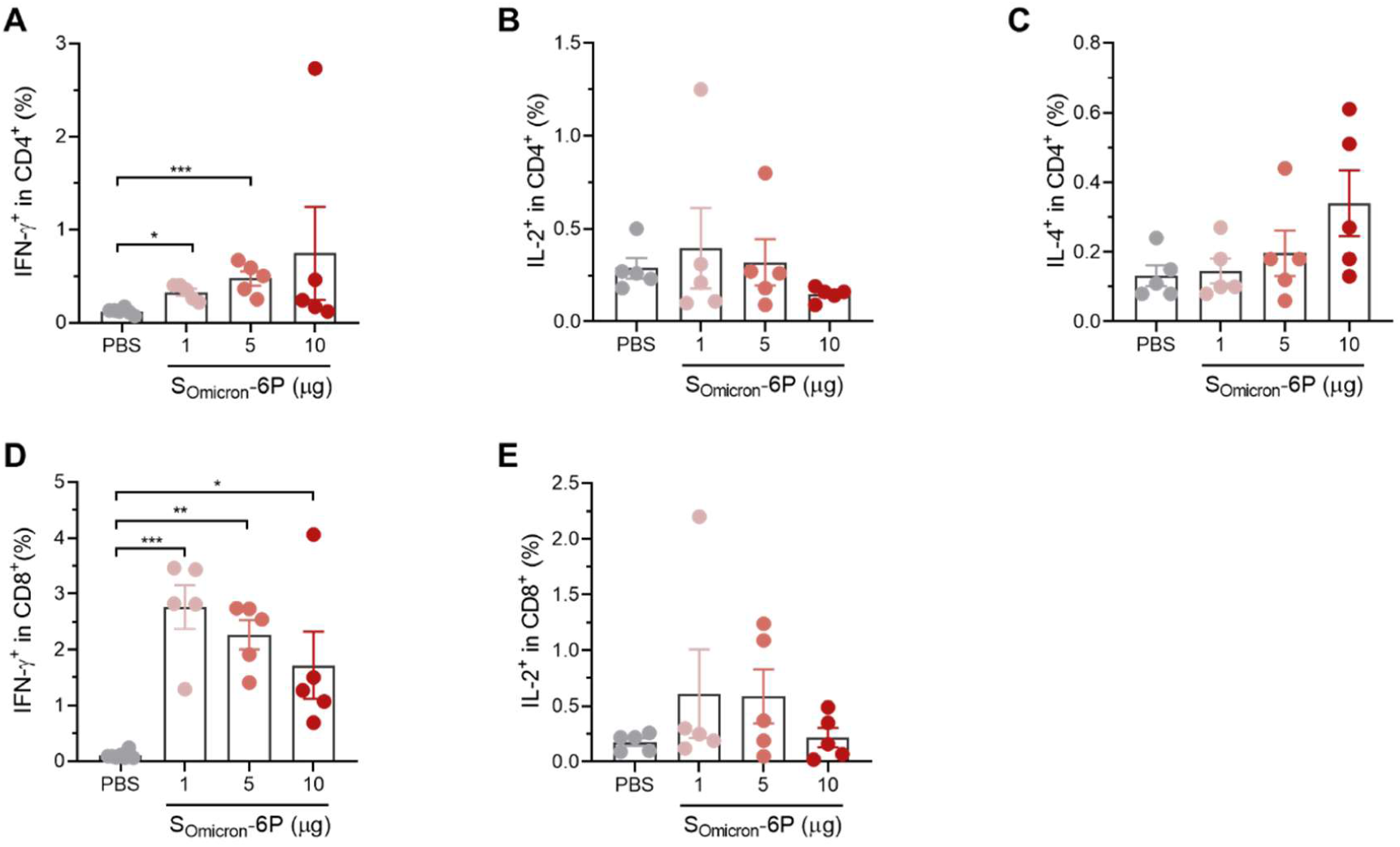
T Cell Intracellular-Cytokine Analysis of SOmicron-6P Immunized Mice, Related to Figure 2. Splenocytes of mice receiving different immunizations were ex vivo re-stimulated with full-length S peptide mix or cell culture medium. Flow cytometry analysis of the percentages of IFN-γ^+^, IL-2^+^, and IL-4^+^among CD4^+^ and CD8^+^ T cells. (A-C) Flow cytometry analysis of the percentages of (A) IFN-γ^+^, (B) IL-2^+^, and (C) IL-4^+^among CD4^+^ T cells. (D-E) The percentages of (D) IFN-γ^+^, and (E) IL-2^+^ among CD8^+^ T cells. Data are shown as mean ± SEM. Significance was calculated using one-way ANOVA with multiple comparisons tests (*p < 0.05, **p < 0.01, ***p < 0.001)

**Figure S9.**
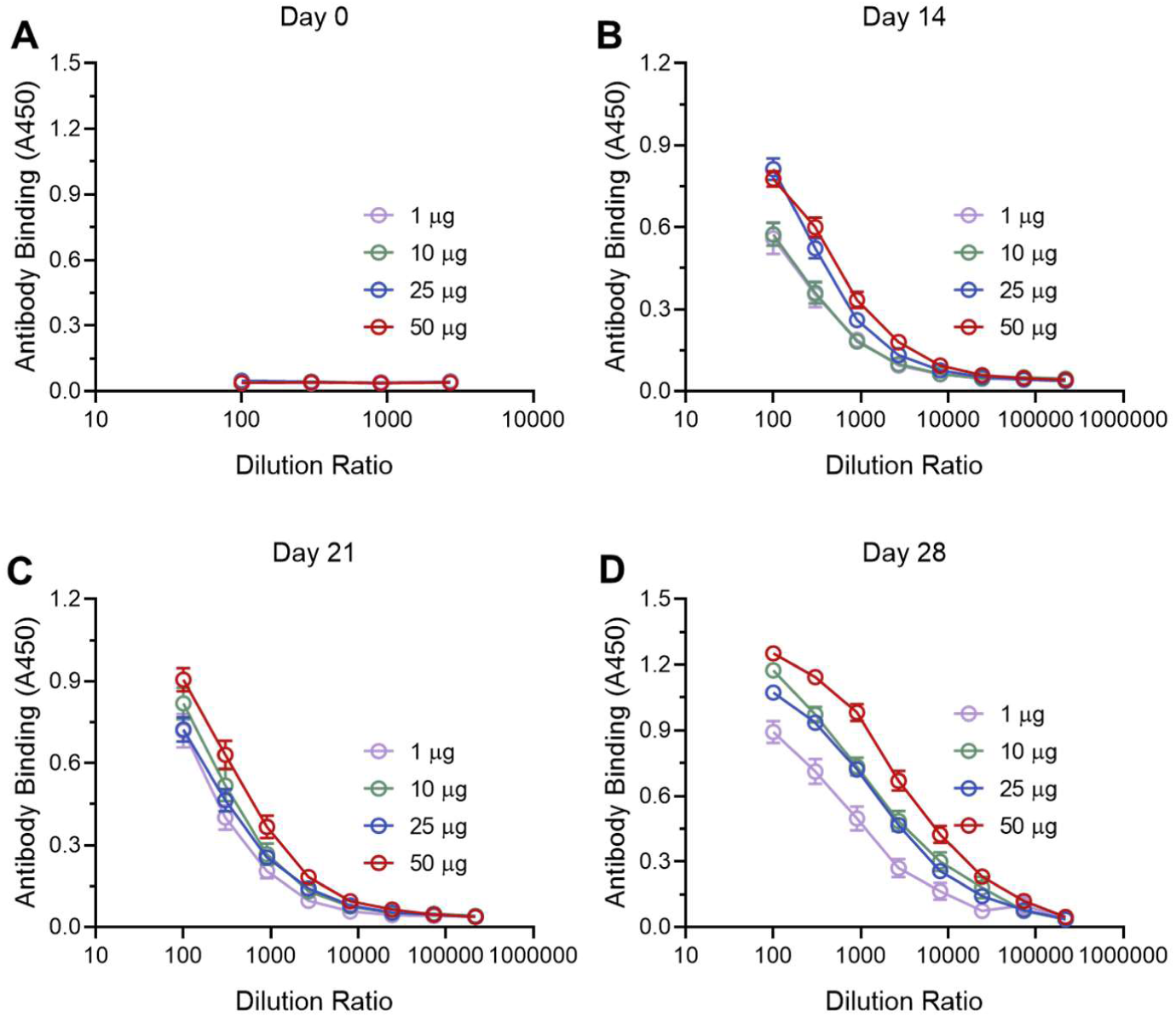
S_Omicron_-6P Elicited Binding Antibodies in Hamsters, Related to Figure 3. (A-D) ELISA binding curves of S_Omicron_-6P induced antibodies in hamster sera on (A) day 0, (B) day 14, (C) day 21, and (D) day 28.

**Figure S10.**
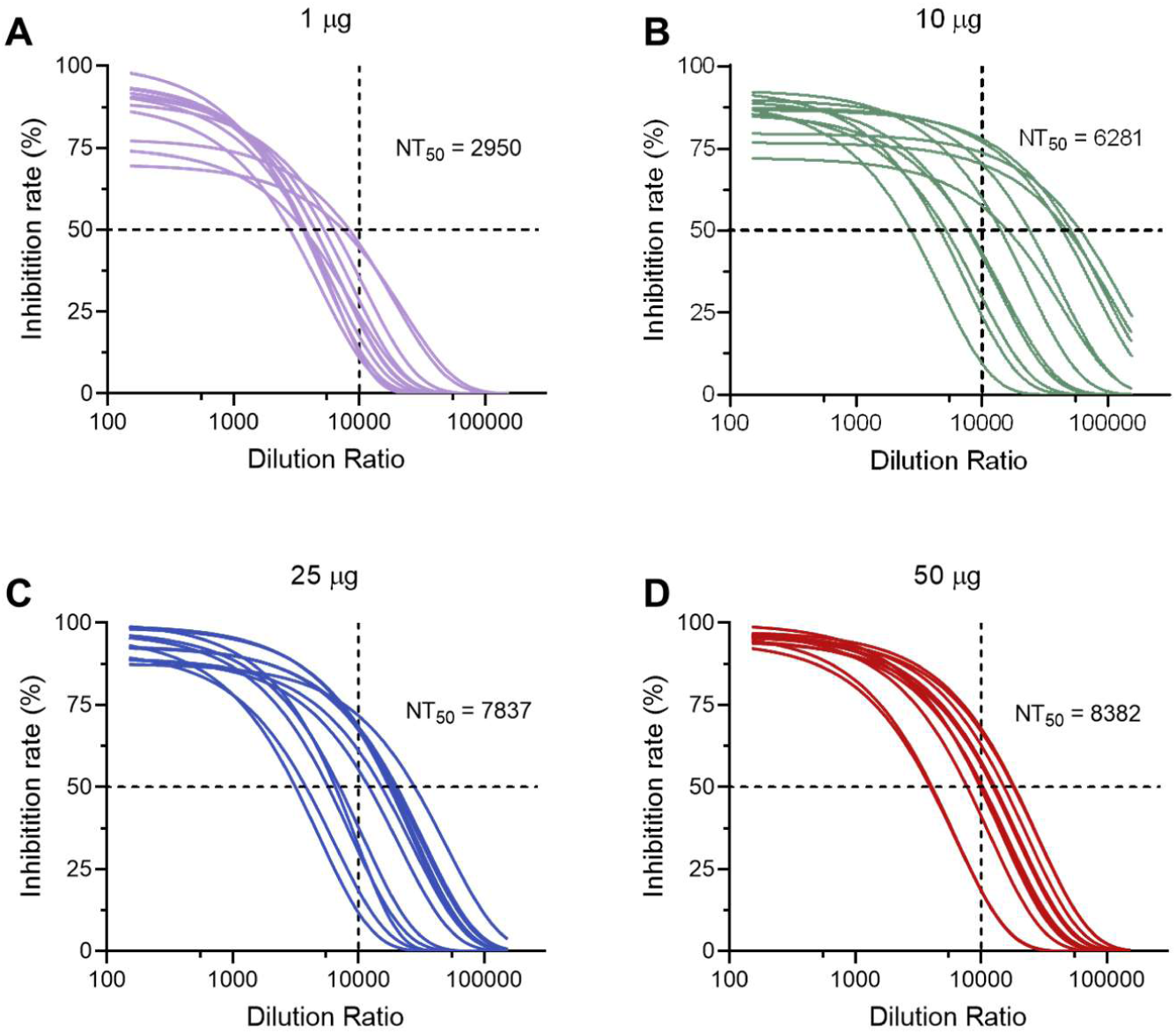
S_Omicron_-6P Induced High Levels of nAbs Against SARS-CoV-2 Omicron Variant in Hamsters, Related to Figure 3. (A-D) Neutralization curves of (A)1, (B) 10, (C) 25, and (D) 50 μg S_Omicron_-6P induced antibodies against pseudotyped and replication-deficient SARS-CoV-2 Omicron at 1 week after second vaccination.

**Figure S11.**
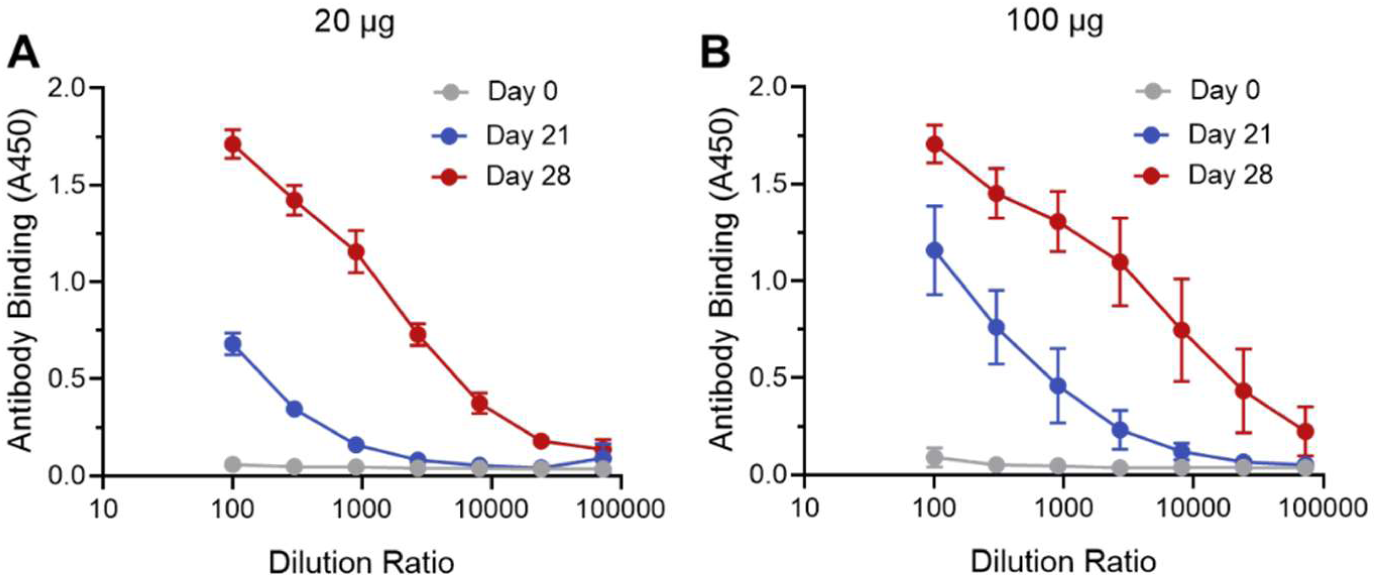
S_Omicron_-6P Elicited Binding Antibodies in Macaques, Related to Figure 4. (A-B) ELISA binding curves of (A) 20 and (B) 100 μg S_Omicron_-6P induced antibodies in macaque sera on day 0, 21, 28 after the first immunization.

**Figure S12.**
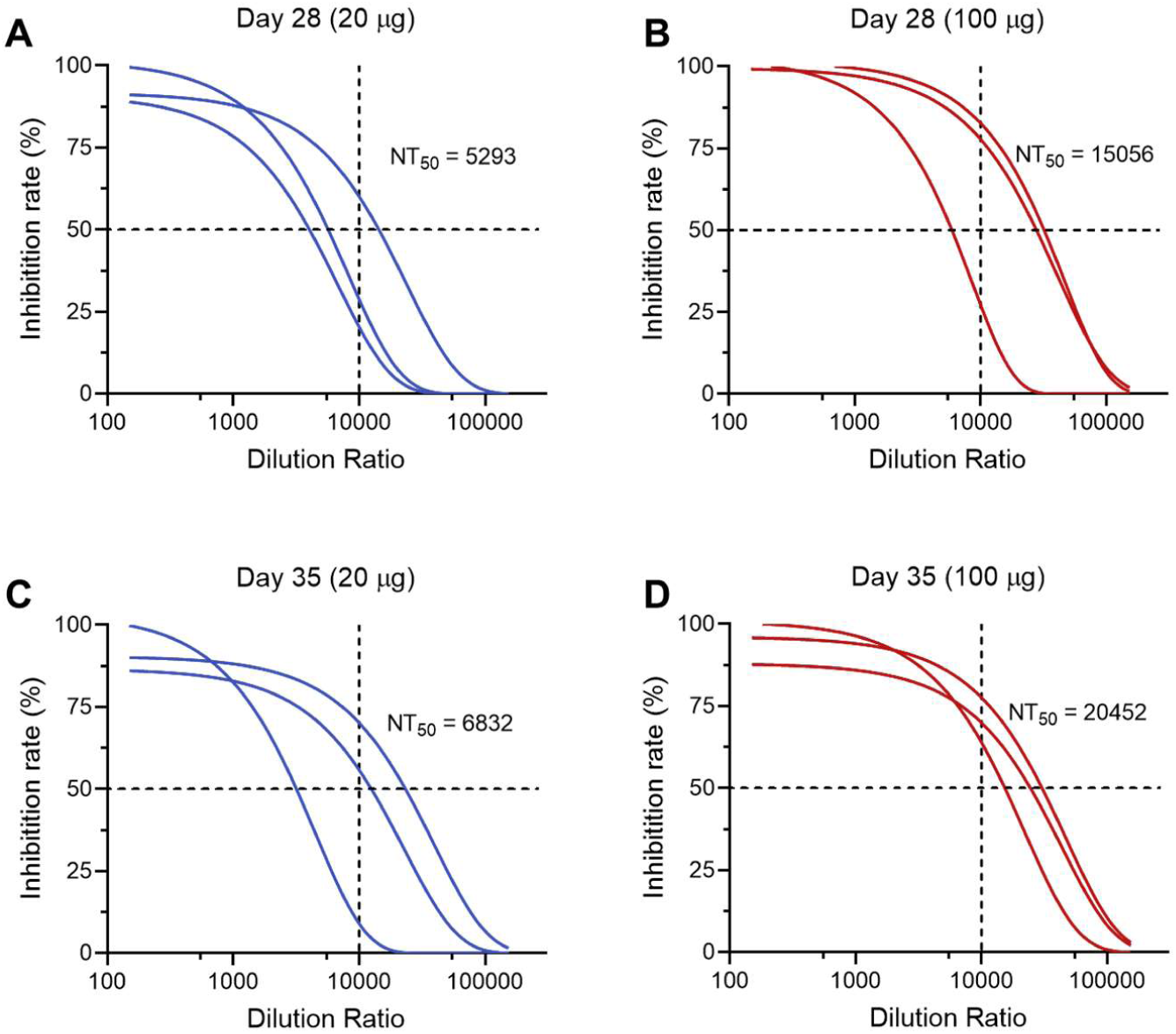
S_Omicron_-6P Induced High Levels of nAbs Against SARS-CoV-2 Omicron Variant in Macaques, Related to Figure 4. (A-D) Neutralization curves of S_Omicron_-6P induced antibodies against pseudotyped and replication-deficient SARS-CoV-2 Omicron (A and B) 1 week and (C and D) 2 weeks after the second vaccination.

## Notes

### Competing Interest Statement

The authors have declared no competing interest.

